# Bayesian Nonparametric Models Characterize Instantaneous Strategies in a Competitive Dynamic Game

**DOI:** 10.1101/385195

**Authors:** Kelsey R. McDonald, William F. Broderick, Scott A. Huettel, John M. Pearson

## Abstract

Previous approaches to investigating strategic social interaction in game theory have predominantly used games with clearly-defined turns and limited choices. However, most real-world social behaviors involve dynamic, coevolving decisions by interacting agents, which pose challenges for creating tractable models of behavior. Here, using a competitive game in which human participants control the dynamics of an on-screen avatar against either another human or a computer opponent, we show that it is possible to quantify the dynamic coupling between agents using nonparametric models. We use Gaussian Processes to model the joint distributions of players’ actions and identities (human or computer) as a function of game state. Borrowing from a reinforcement learning framework, we successfully approximated both the policy and the value functions used by each human player in this competitive context. This approach offers a natural set of metrics for facilitating analysis at multiple timescales and suggests new classes of tractable paradigms for assessing human behavior.

---

Over the last fifteen years, game theory has been foundational in establishing cognitive and biological mechanisms of strategic decision making [1–4]. Paradigms like Matching Pennies, the Trust/Ultimatum Games, and Prisoner’s Dilemma have used simple choices in highly standardized contexts to rigorously characterize the psychological processes underlying social concepts such as trust, altruism, and inequity aversion. The growing adoption of paradigms from game theory has yielded key insights into social decision-making in humans [4–8] and animals [9–11]. These game theory paradigms draw upon a vast literature detailing how rational players would behave [1,4,12–14], yet studies comparing human behavior to these normative solutions have found that humans often violate rational predictions [1,4,12,15].

While a central aim of game theory is to describe how people *should* make decisions, describing how humans *actually* make decisions is of particular interest to social scientists. Indeed, many of the features that have made game theory paradigms analytically attractive—discrete choices, turn-taking, known payouts—are abstractions away from real-world social interactions. For instance, when buyers haggle over the price of a good, they respond to one another in real time, using a combination of nonverbal cues, strategic planning, perspective taking, and value judgment. Their continuous, dynamic interaction thus forms a challenge to any computational framework for the study of social decisions [4,16,17]. Moreover, while game theory has proven highly successful in analyzing various sorts of equilibria players might settle into, considerably less is known about the processes by which these equilibria are reached [18,19], As a result, it is desirable to develop analytical tools capable of quantifying strategic dynamics while maintaining the mathematical rigor that has made game theory such a productive framework.

Here, we introduce a computational modeling framework that borrows heavily from recent advances in reinforcement learning [20–27] and nonparametric Bayesian modeling [28–30] to capture these social dynamics. Our approach produces models of behavior that are both flexible enough to capture the variability present in a continuously evolving strategic setting and powerful enough to quantify strategic differences across participants, trials, and even individual moments within trials. Our testbed for these ideas is a competitive task in which human participants played against both a human opponent and a computer opponent in a real-time, movement-based game. This paradigm generates a rich complexity in individuals’ behavior that can be succinctly described by individualized, instantaneous policy and value functions, facilitating analysis at multiple timescales of interest. This approach serves to quantify complex interactions between multiple agents in a parsimonious manner and so suggests new classes of tractable paradigms for studying human behavior and strategic decision making.

## Results

### Penalty Shot Task

We adapted a zero-sum dynamic control task, inspired by a penalty shot in hockey [17], The task was viewed on a computer screen and played by two players: an experimental participant (n = 82) who controlled an on-screen circle, or “puck,” and another long-term participant who controlled an onscreen bar, acting as the “goalie,” Hereafter, we will refer to these players as the participant and the opponent, respectively. The puck began each trial at the left of the screen and moved rightward at a constant horizontal speed. The task of the participant was to score by crossing a goal line located at the right end of the screen behind the opponent. The opponent’s task was to block the puck from reaching the goal line. Each player moved his or her avatar using a joystick. Both players were only able to control the vertical velocities of their respective avatars, though the puck and bar had distinct game physics (see Methods and Supplement Methods 1), See Fig 1 for task progression and sample trajectories, as well as the Supplement Video 1 for a video demonstrating real game play.

**Figure 1:**
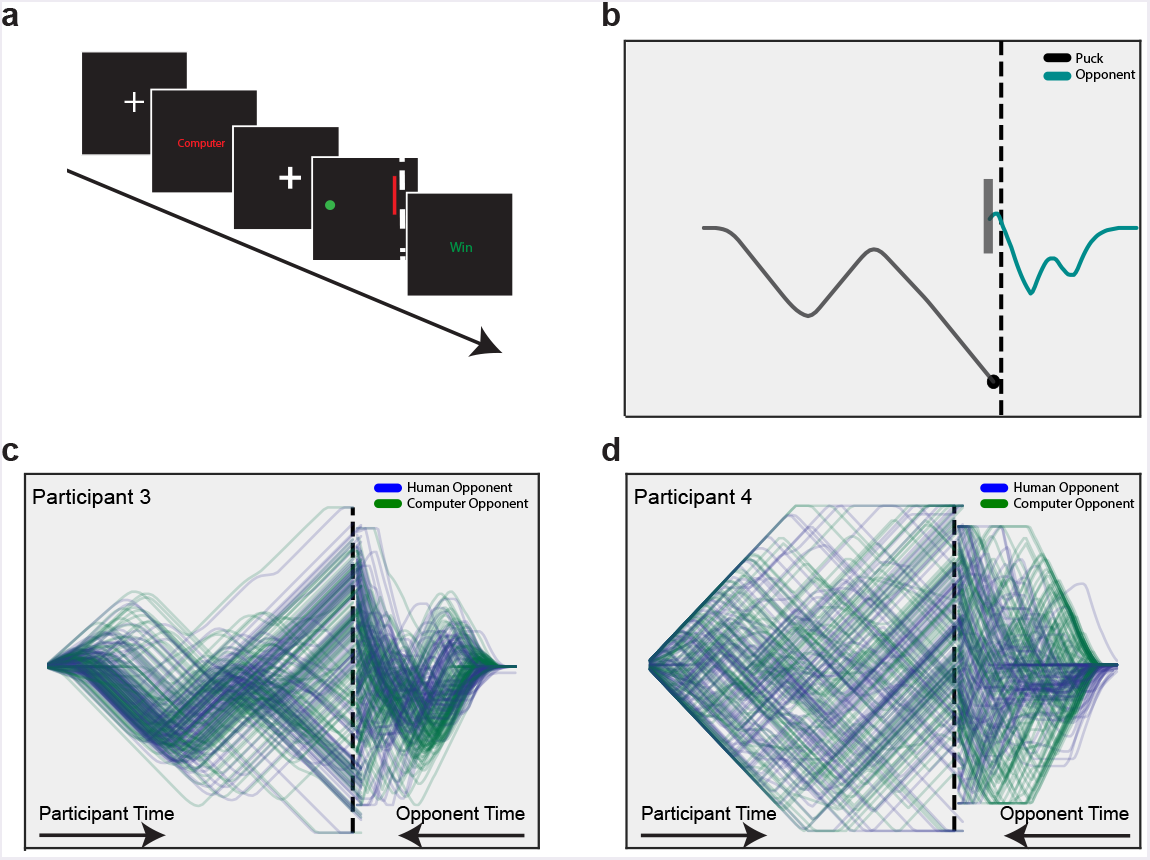
Strategic heterogeneity in dynamic decision-making. A: Task progression: Following a jittered fixation cue, text indicated the identity of the opponent on the upcoming trial for 2 seconds. Play commenced after a variable delay during which the screen displayed a fixation cue. At the conclusion of each trial, which lasted roughly 1.5 seconds, colored text indicated the winner (green “Win” if the participant won; red “Loss” if the participant lost) for 1.5 seconds. B: Game play on a single trial. The puck moves from left to right at constant horizontal velocity. The bar was only allowed to move vertically, but is depicted as moving from the right side of the screen inward toward the goal line for visualization purposes. C and D: All of the trajectories for Participant 3 (C) and Participant 4 (D), demonstrating the heterogeneity observed across participants. Note variability in both on screen positions’ visited and trajectory shape: Participant 3 is much more consistent in game play, while Participant 4 was more variable. Trials played against the human opponent are displayed in blue. Trials played against the computer opponent are in green.

Participants played the penalty shot task in an fMRI scanner. Here, we report only the behavioral data from this experiment. On half of the trials, the experimental participant played against the human opponent, located outside the scanner. On the other half of trials, the participant played against a computer-controlled opponent. The computer opponent followed a “track-then-guess” heuristic in which it attempted to match the puck’s vertical position (with a variable reaction time) before randomly choosing a direction to move at maximal speed near the end of the trial. This choice was motivated not only by pilot data that showed such a strategy was difficult for participants to exploit, but also by past work analyzing the anticipatory strategies of goalkeepers [31–33], Opponent identity was randomly selected on each trial. Our task was incentive-compatible: both the experimental participant and the human opponent were rewarded in monetary bonuses that were dependent on how frequently each player won.

As expected, participants exhibited considerable variability in game play. Figure 1 shows all trajectories for a representative pair of subjects. Clearly, a salient feature of our paradigm is its accommodation of widely varying individual strategies, (As a result, for each of our main analyses, we only display findings for a subset of participants. Plots for all representative subjects for all analyses are available in Supplementary Figures 1-8, Trajectories varied widely both within and across participants, despite the fact that players each only had one continuous degree of freedom (position along the y-axis). For example, Participant 3 (Figure 1C) demonstrated highly stereotyped play, with most trials exhibiting a “down-up-guess” approach. By contrast, Participant 4’s (Figure 1D) trajectories were dispersed throughout the screen, perhaps resulting in less predictable play. Participants also experienced highly variable win rates, which ranged from 43-76% (against human: 34-83%; computer 42-73%).

### Gaussian Process Models

Our observed data for each trial were movement trajectories for the puck and the bar, each spanning approximately 1.5 seconds (94–96 discrete time points). While it is possible to model these time series directly [17], we observed that for many participants, puck trajectories could be characterized as comprising a series of straight-line segments of maximal or near-maximal velocity separated by change points (Fig 2A). That is, we could redefine the decision available to the participant at each moment as whether or not to switch direction. This transforms a time series modeling problem into a more tractable change point prediction problem, for which our predictors are a small number of game state variables.

Viewed through the lens of reinforcement learning, the decision of whether to switch direction at time *t* is an action, *a_t_*, and the probability of this action given a state of the world *s_t_* is given by the policy function: Π(*a_t_,s_t_,ω*) = *p*(*a_t_*|*s_t_,ω*), where we let *s_t_* denote a vector of predictors at each time point *ω* is a binary variable indicating the opponent’s identity (computer = 0, human = 1) [20], In principle, both states and actions can be continuous, though in practice, they are often discretized [20,34], In our case, we define the action space as a single binary variable, with 1 indicating a change in direction and a 0 indicating continuation along the current trajectory. However, the state *s* remains continuous and includes a total of 7 predictor variables: the *x* and *y* positions of the puck, the *y* position of the bar, their respective vertical velocities, the time since the occurrence of the last change point (normalized to 1 by dividing by total trial length), and an opponent experience variable that ranged from 0 (first trial) to 1 (last trial) that was specific to each opponent and reflected potential strategic adaptation over the course of the experiment. Finally, we simplify our notation, defining *π*(*s_t_,ω*) = *p*(*a_t_* = |*s_t_,ω*). Because our input space is of moderate dimension, a model for *π*(*s, ω*) will be a continuous function of s instead of a large matrix, as it would be for a model with a discrete state space. Our contribution is to show that nonparametric methods allow us to address the challenge of modeling *π* using only sparsely sampled data.

Our decision to model change point probabilities as a function only of states and opponents means that the data at each time are independent of each other given these variables. Thus, our approach is also equivalent to a binary classification problem. Binary classification is well-studied, with many methods available, including logistic regression, support vector machines, and neural networks [35]. Our selection of model was guided by three requirements: First, the model should be flexible enough to capture the rich diversity of player behavior. Second, the model should appropriately handle a small number of change points (≈ 4.6%) with an input space of moderate dimension. And third, the model should avoid overfitting while providing a principled estimate of uncertainty. For these reasons, we fit each participant’s data using a Gaussian Process (GP) classification model,

A Gaussian Process (GP) is a distribution over functions, GPs are widely used in spatial and time series modeling for their combination of flexibility and ability to generalize from even modest data [28,36], In the same way that a sample from a normal distribution is a real number and a sample from a Bernoulli distribution is a binary variable, a sample from a GP is an entire function (e.g., a univariate time series (*d* = 1) or spatial density (*d* = 2)). Gaussian Processes have the advantage of providing a principled, Bayesian measure of uncertainty over functions while remaining resistant to overfitting and generalizing to unseen data [28]. They are also equivalent to single-layer, fully-connected, infinitely wide neural networks, and have been shown to outperform neural networks in avoiding overfitting on small to moderate datasets [37–39]. Moreover, they are the method of choice when modeling time courses based on sparse or irregularly-sampled data [40,41]. Thus, GPs offer competitive modeling performance with the added benefits of uncertainty estimation and differentiability.

More formally, a GP *f* is defined by a mean function *m*(*x*) (usually assumed to be 0 *a priori*) and a covariance function *k*(*x,x′*) that defines the correlation between values of *f* at different input points [28]:

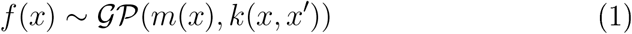

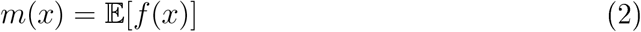

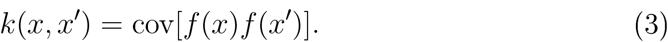

By definition, the joint distribution of the observed data set 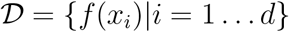 is multivariate normal with dimension *d*, mean *μ_i_* = *m*(*x_i_*), and covariance Σ_*ij*_ = *k*(*x_i_,x_j_*).

As stated above, we chose to model players’ policies via a GP classification model that attempted to predict an upcoming change in the puck’ direction from the current state *s* and opponent identity *ω*. Following standard techniques [28,42], we assumed that binary change point observations *a_i_* were Bernoulli distributed according to the policy *π*(*s,ω*) and that the policy itself was related to an underlying GP:

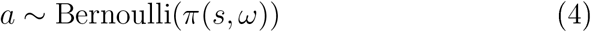

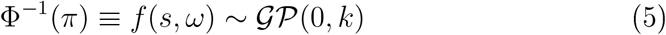

where Φ^−1^ is the inverse cumulative normal distribution (also called the probit or quantile function) and 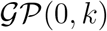 is a GP prior on *f* with mean 0 and kernel function *k*. Because we assume that *f* is a smooth function of its inputs, we choose the common radial basis function (EBF) kernel [28]:

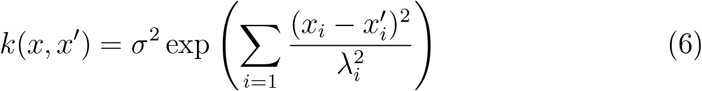

with *i* indexing input variables and *σ_i_* and *λ_i_* hvperparameters setting the overall magnitude of the covariance and the length scale of correlations along each dimension, respectively. Here, *x* includes both s and *ω*. Even though *ω* is a discrete parameter, we approximate it as a continuous variable, as is often done in Bayesian modeling using GPs [43].

**Figure 2:**
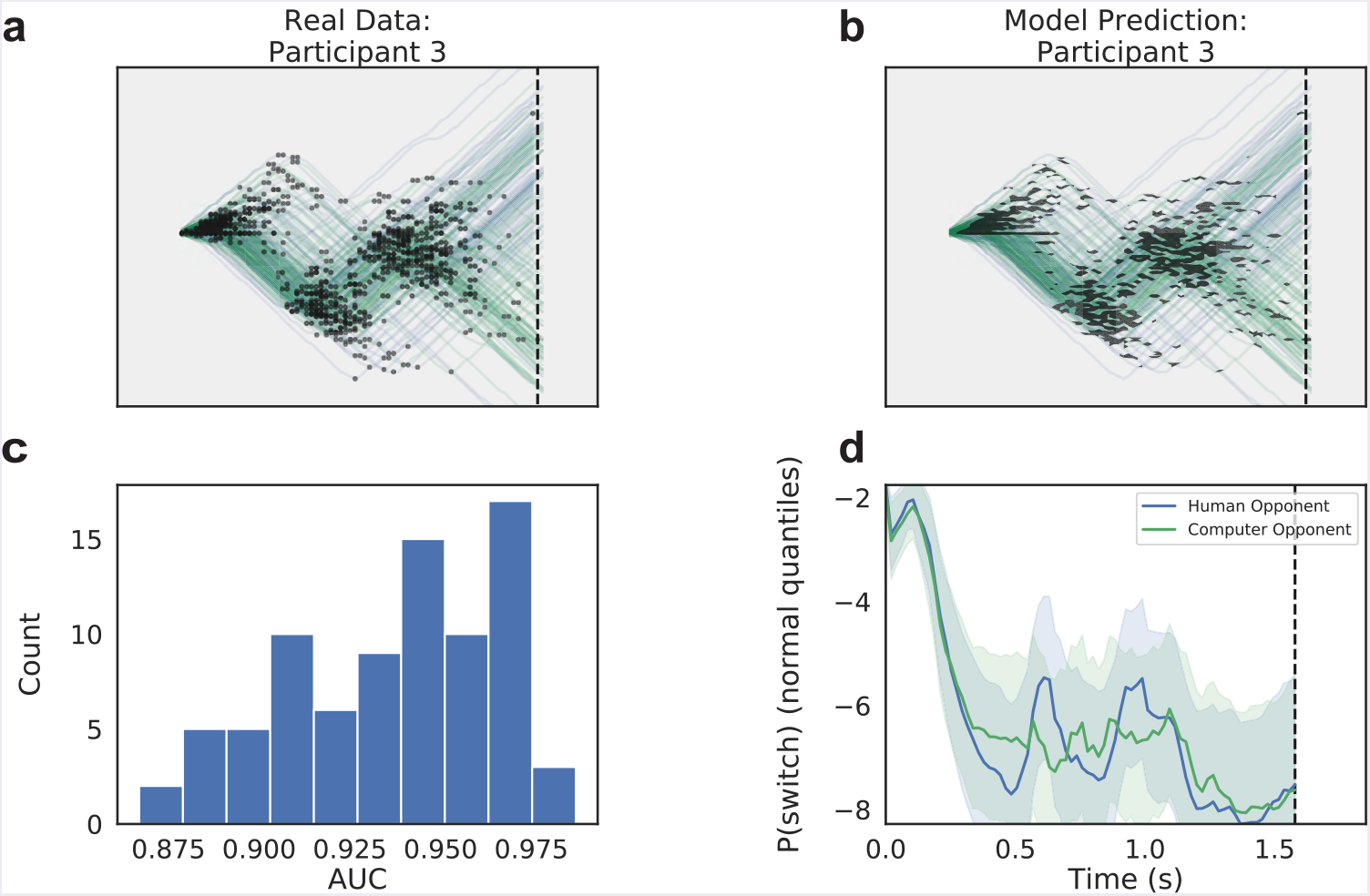
A change point classification model captures variability in player strategy. A: Observed data from a single subject (Participant 3) in the penalty shot task. Blue trajectories correspond to trials played against the human opponent. Green trajectories are from trials played against the computer. Black dots represent change points, or switches in joystick direction by the participant. B: Same trajectories from A, but overlaid with black shaded regions indicating locations in which the model-predicted probability of a change points exceeded the participant’s base rate. C: Histogram of participants’ area under the curve (AUC) scores on held-out data (20% of each participant’s dataset). D: Probability of a change point as a function of time, averaged across trials, for Participant 3. Shaded regions indicate 95% Bayesian credible intervals. Probabilities are shown and averaged in quantiles (z) of the normal distribution. Blue indicates trials against the human opponent, green against the computer.

We found that our GP classification model accurately captured the diverse patterns present in participants’ data (Fig 2A,B), That is, the model had a higher probability of predicting a change point in regions of the screen where change points actually occurred. This is a direct result both of the nonparametric nature of the GP—the model adapts its complexity to the data—as well as the smoothing effects of the prior, Held-out test data from each participant yielded a median area under the curve (AUC) score of 94% (Fig 2C), For comparison, we also fit a logistic regression to each subject, but for no subject did it outperform our GP model (see Supplement Figure 9).

### Disentangling identity and context effects in play

We next wanted to investigate how player strategy differed based on the identity of the opponent (human or computer). As Figure 2D illustrates some participants evinced minimal differences in switch probability between the two opponents. However, this contrast elides an important distinction between what might be termed “opponent identity effects” and “opponent context effects”. That is, we might ask whether observed differences in switch probability between the two opponents are due to intrinsic differences in the way participants perceive each opponent or the fact that each opponent simply plays a different strategy. In typical social games, these effects are all but impossible to disentangle, but because we model the joint distribution of both states and opponent identity, *f*(*s,ω*), we can perform the following “counterfaetual” experiment: For every state s visited in play against the computer (*ω* = 0), we can ask how *f*(*s*,0) compares to *f*(*s,* 1), This is equivalent to freezing game play at a single moment, switching the identity of the opponent while holding all other variables fixed, and asking how play in the next instant differs. Such a pure identity effect quantifies how much participants’ strategies would differ between human and computer opponents who used the same strategy.

In fact, the observed contrast between the two curves in Figure 2D can be fully decomposed into an effect due to opponent identity and an effect due to differences in the distributions of visited states (see Methods), As indicated in Figure 3A, the observed contrast plotted in Figure 2D corresponds to the difference along the diagonal, while the identity and context effects correspond to differences taken along the vertical and horizontal directions, respectively. Figures 3B and C illustrate this decomposition for two representative participants. These figures show both the observed contrast (difference between the two curves in Figure 2D and its constituent pieces due to opponent identity and context. While the latter are typically larger, indicating a predominance of game state effects on switch probability, there is considerable heterogeneity across both participants and time in trial. Figure 3D illustrates this by considering the average identity effects for each participant during the early and late stages of each trial. There, a positive value indicates higher switch probability for a human opponent, while a negative value indicates higher switch probability against the computer. While some participants consistently exhibit higher switch probabilities against the human opponent (upper right) or against the computer (lower left), others switch more against one early and the other late (upper left, lower right). Thus, players can be distinguished not only by which opponent elicits more switching behavior, but also by the periods of the trial in which these tendencies occur.

### Sensitivity to opponent actions differs between human and computer play

We next sought to quantify how much participants’ switching behavior changed as a function of the opponent’s *actions*. Because our change point policy model is based on a smooth Gaussian Process, we could naturally quantify this sensitivity using gradients of the GP *f* = Φ^−1^(*π*) with respect to the opponent’s position and velocity (see Methods), We then used these gradients to define a moment-by-moment *sensitivity index*. Since the gradients of the GP measure the degree to which small changes in the current game state affect the participant’s probability of changing course, gradients with respect to the opponent’s position and velocity capture the degree to which the participant’s current behavior is sensitive to the opponent’s actions at each time.

As Fig 4A illustrates, like the probability of switching, sensitivity to opponent action varied throughout the trial. Even when opponent effects are disambiguated from context (Fig 4B, C), clear effects remain for most participants, In fact, repeating the analysis of Fig 3D for our sensitivity metric reveals an even greater diversity in strategic variation (Fig 4D), Just as we saw with the behavioral trajectory and the likelihood of switching, participants exhibit significant variation in both the magnitude and timing of opponent-related differences in sensitivity.

**Figure 3:**
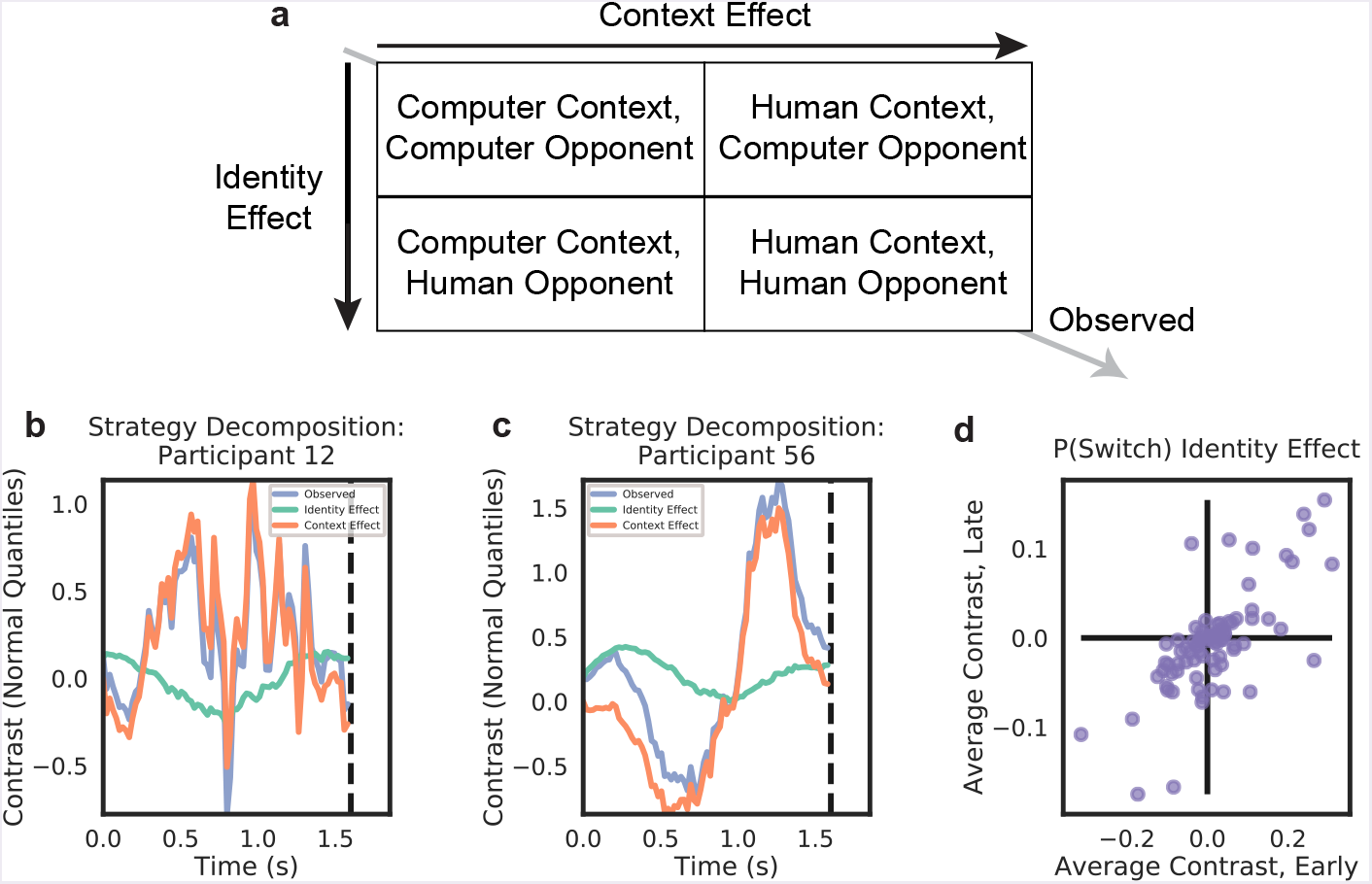
Disentangling identity and context effects. A. Schematic of the identity versus context decomposition. Differences in the expected values of model variables between human and computer opponents can be decomposed into a sum of identity and context effects (see Methods). B. Decomposition as a function of time in trial for Participant 12. The difference between human and computer switch probabilities (in quantiles; purple) is the sum of opponent (green) and context (orange) effects. C. Same decomposition as in B, for Participant 56. D. Population variability in opponent effect. Scatter plot of trial-averaged switch probabilities for the first and second half of the trial for each participant. Participants in the upper right consistently switch more against the human opponent, participants in the lower left against the computer. Participants in the other two quadrants switch more frequently against one opponent in the early half of the trial and reverse this behavior in the latter half.

### Sensitivity metrics characterize behavior across multiple time scales

The sensitivity metric defined above represents a particular moment-by-moment measure of the degree to which one player (the participant) is coupled to the actions of the other (the opponent). Based on our prior expectation, we chose a combination of sensitivities to opponent position and velocity, but other combinations are equally plausible. In fact, one could define a sensitivity metric to each input variable individually. Here, we show that such an approach is not only feasible, it produces a principled characterization of participants’ behavior across multiple timescales. Indeed, when aggregated at the participant level, these indices fully characterize the policy model.

**Figure 4:**
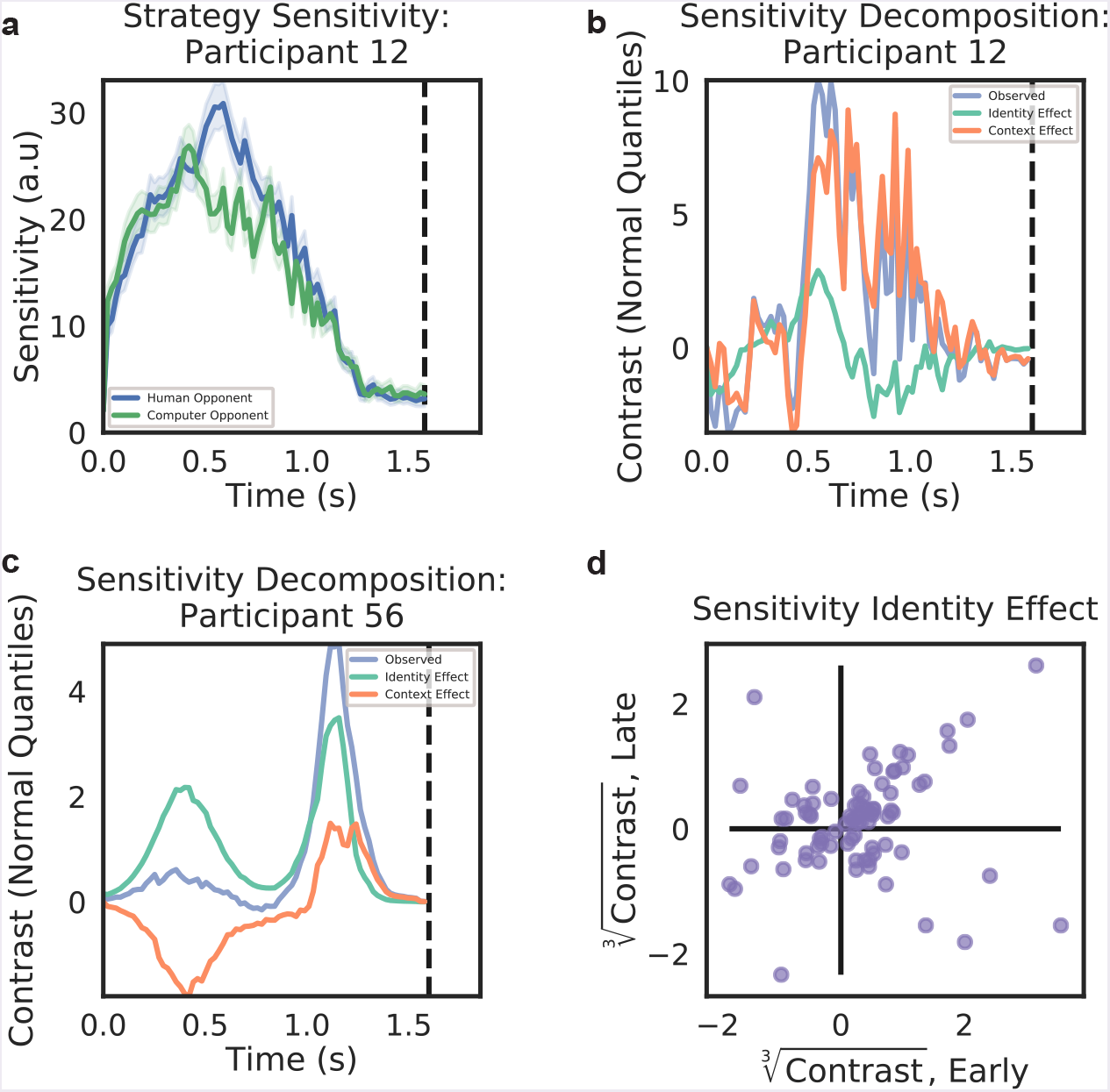
A gradient-based measure of sensitivity to opponent action reveals distinct information about player strategies. A: Observed sensitivity to opponent actions in both opponent conditions, for Participant 12. Shaded regions indicate 95% credible intervals. Blue line and shaded region correspond to the human opponent condition, green to the computer opponent. B: Same participant as in A, with the observed contrast decomposed into identity and context effects, similar to Fig 3B. C: Same as B, but for Participant 56. D: Population variability in sensitivity to opponent actions. Conventions are as in Fig 3D, but sensitivities have been cube-root transformed for visualization purposes.

By analogy with the approach described in the last section, we defined one sensitivity for each input variable, equal to the square of the gradient along each input direction (see Methods). This yielded eight new sensitivity indices (seven for state plus one for opponent identity) in addition to the opponent action sensitivity defined above. However, our previous index can be defined in terms of these new indices, so there are only eight unique values in the set. The most important feature of these new indices is that, like the policy, they are defined moment-bv-moment, but can be aggregated across multiple levels of granularity, including trial and participant averages. We have already taken advantage of this in Fig 3D and Fig 4D to illustrate variance in switch probability and sensitivity across our population, but one can also approach this more systematically. As in classic analysis of variance (ANOVA), we can consider each index value at each data point as the sum of three terms: a participant-level mean, a trial-level offset from this mean, and a residual specific to the data point. Likewise, we can use the data to estimate variances within trial (residual), across trials, and across our participant population. As in ANOVA, the sum of these variances, appropriately weighted, equals the total variance in the data. Normalizing by this total variance yields a set of three positive terms that sums to 1:

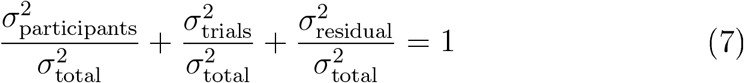

Points following Eq 7 lie inside the triangle shown in Fig 5A, Points close to the lower left corner represent variables whose variance is almost entirely accounted for at the within-trial level, while points nearer the lower right and apex represent variables with relatively more variance across trials and participants, respectively. Unsurprisingly, points are skewed toward the lower left, since most variance is at the time point level. This stems from the fact that points early and late in the trial are much less like each other than points near the same time across trials. Nonetheless, the actual strategy, as summarized by the baseline probability of switching, exhibits a sizable variance across participants, indicating that the differences in change point frequency apparent in Fig 1C and D are relatively more “trait-like” than our sensitivity measures. Perhaps surprisingly, while the sensitivities to opponent position and velocity are almost entirely characterized by within-trial variance, the aggregated metric defining opponent action sensitivity has more variance accounted for within participants, suggesting it to be more trait-like than the opponent position and velocity sensitivities alone.

**Figure 5:**
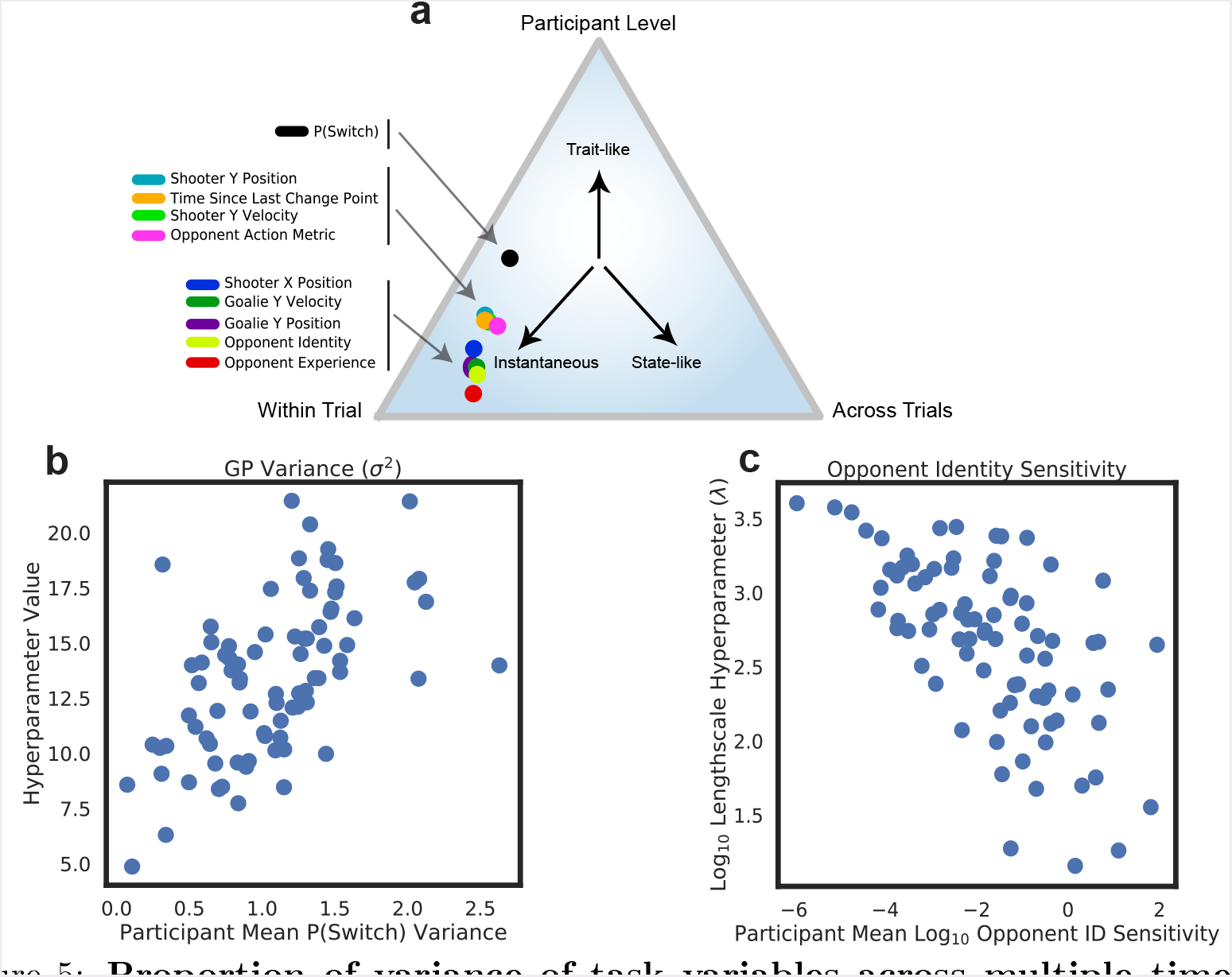
Proportion of variance of task variables across multiple time scales. A: Variance decomposition of model metrics. Points inside the triangle represent different allocations of variance within-trial, across-trial, and across-participant. Points near the apex represent more “trait-like” variables, while variables near the lower left are more “state-like.” Our metrics, place most of their variance within trial, since data points vary more across time than across participants. B: Relationship between mean probability of switching and GP noise parameter across participants. Each dot represents one participant. C: Relationship between sensitivity to opponent identity and GP opponent identity hyperparameter.

Finally, we note an important relationship between participant-level sensitivities and model hyperparameters. Because Gaussian Processes, like Gaussian distributions, are completely characterized by their mean and covariance, summaries of a Gaussian Process taken over the data can only be functions of the hyperparameters that define the mean and covariance. That is, we expect on mathematical grounds that our sensitivities, when averaged across an entire participant’s data, should be simply related to the model’s hyperparameters (see Supplementary Note 2). Fig 5B and C illustrate this relationship for two hyperparameter-sensitivity pairs. Fig 5B shows that the noise parameter of our Gaussian Process model, *σ*^2^ is indeed correlated with the variance in probability of switching across time points for each participant (*R* = 0.56, *t* = 6.09, *p* < 0.0001), Likewise, Fig 5C shows that the logged hv-perparameter controlling opponent identity effects in the GP correlates negatively with each participant’s sensitivity to the same variable (*R* = −0.64, *t* = 7.43, *p* < 0.0001), In both cases, this is exactly what we expect: the noise hvperparameter for a classification model is related to the variance in its predictions, while low sensitivities correspond to long correlation length scales. Thus, our gradient-based sensitivity metrics straightforwardly and naturally extend GP hyperparameters to the time point level, providing a principled characterization of strategy suitable for analysis at multiple timescales.

### Action Value Model

We have shown that we can use nonparametric methods to estimate the policy participants use when playing a dynamic, strategic game. Yet this analysis says nothing about how effective these policies are. So how do participants’ choices at each moment translate to wins and losses? To answer this, we separately modeled each participant’s action value *Q_π_*(*a*|*s,ω*): the expected value of taking action *a* in state s against opponent *ω* and playing according to policy *π* thereafter. As indicated by notation, this value is policy-dependent. That is, each policy *π* uniquely determines a value function *Q_π_*. In typical reinforcement learning models, policies are likewise dependent on action values: Given action values, Q, policies choose actions based on a softmax function or other rule [20], Thus, there is a mapping in the reverse direction from action values to policies. The Bellman Equation stipulates that for optimal learners, the optimal policy and action values determine one another [20], but this need not hold for nonoptimal learners.

Figure 6A illustrates these concepts. While the optimal policy *π*_*_ and *Q*_*_ are mapped onto each other by the processes of value calculation and action selection, respectively, for non-optimal learners, the observed policy *π_obs_* leads to a value function *Q_obs_*, but softmax action selection based on *Q_obs_* may not be equivalent to the original policy: *π_Q_* ≠ *π_obs_*, so the mappings in Figure 6A are not inverses except for optimal policies. In other words, learners may not necessarily be choosing based on the expected values of their actions. As a result, we took an approach in which the action value function *Q*(*a*|*s,ω*) was modeled independently of *π*: This model took as inputs the instantaneous state, opponent, and observed action at that time and attempted to predict from those data whether the participant subsequently won the trial. We used the same Gaussian Process classification approach as before, only this time predicting the trial outcome and using the participant’s observed action as an additional input.

**Figure 6:**
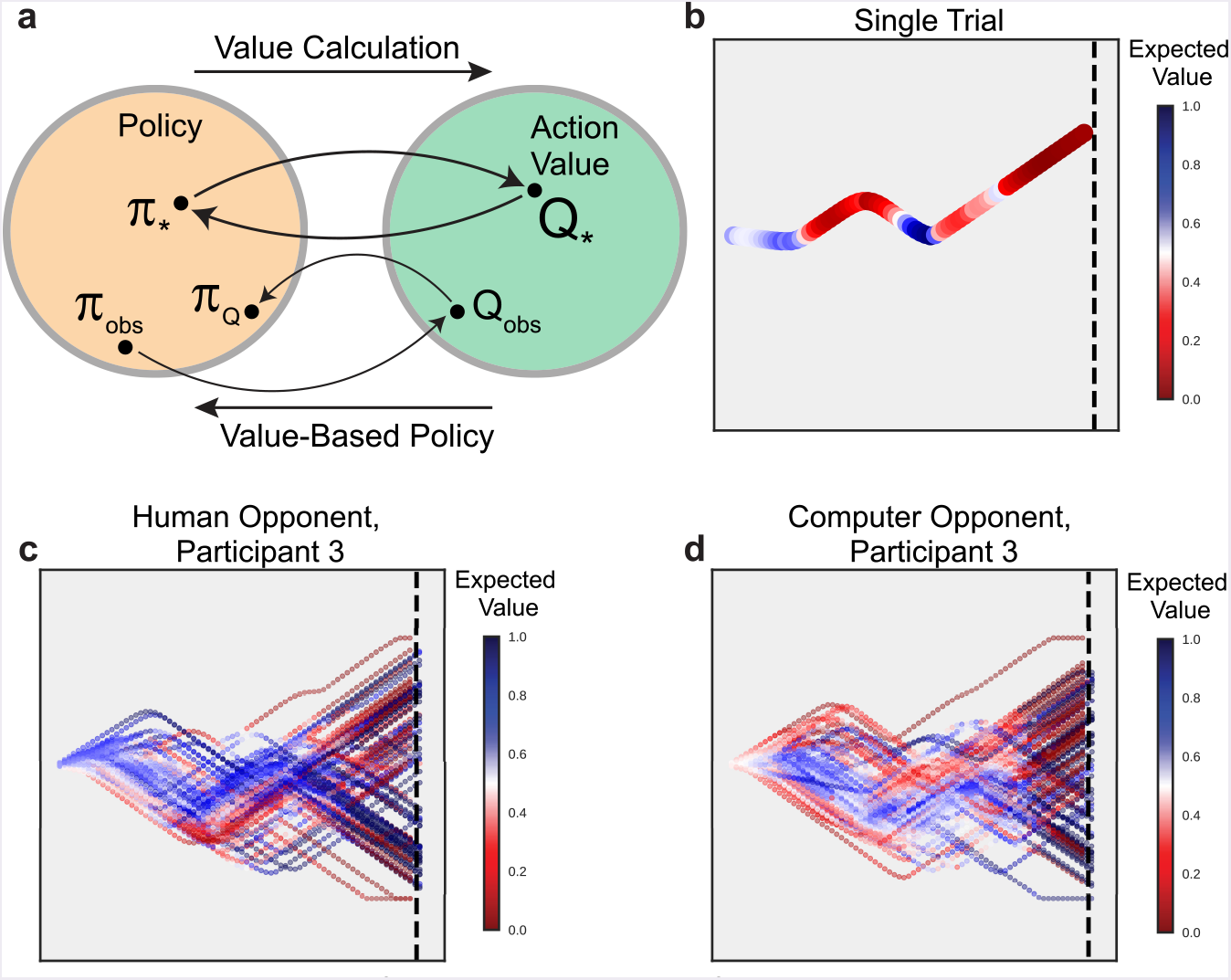
A Gaussian Process action value model captures variability in player efficacy. A: Relationship between policies and action values in reinforcement learning. Each policy determines an action value (rightward arrow). Conversely, a set of action values, coupled with an action selection mechanism like softmax or greedy methods, determines a policy (leftward arrow). For optimal learners, the connection between the optimal policy *π_*_* and its resulting action values *Q*_*_ is given by the Bellman Equation, which states that the leftward and rightward arrows are inverses of one another. For non-optimal agents, however, the observed policy *π_obs_* determines *Q_obs_*, but action selection based on *Q_obs_* may not be the same as *π_obs_*. B: Expected values (win probabilities) at each moment for a single trajectory from one participant. Horizontal and vertical axes correspond to position on the computer screen. Color indicates expected value. C: All trajectories for a single participant against the human opponent. D: Trials against the computer opponent for the same participant as in C. Note the increased intensity of colors late in the trial, after the opponent has made its last move.

The results of this model are shown in Figure 6, As Figure 6B illustrates, there are fluctuations in expected value even within a single trial as players move and counter move. Here, we have plotted the predicted expected value, which is equal to the value function of reinforcement learning: *V_π_*(*s,ω*) = *∑_a_ π*(*a*|*s,ω*)*Qπ*(*a*|*s,ω*), an undiscounted, weighted sum of action values according to their probability under the current policy. Quantifying expected value at the time point level, rather than the trial level, allows us to see how fluctuations in game state impact likelihood of winning. For reference, we fit a logistic regression using the same set of input features and targets. Once again, the GP model outperforms logistic regression for each participant in our cohort (see Supplement Figure 10),

Figures 6C and D show these predictions across all trials for a pair of representative participants. Interestingly, while the types of trajectories generated by Participant 3 in both the human and computer opponent conditions look remarkably similar, expected values for these collections of trials evolve quite differently. Evidently this participant, while playing essentially the same strategy in both cases, experienced much different win rates against the two opponents. In particular, against the computer opponent, we see a more abrupt transition in expected value between the first and second half of the trial. This can be explained as a byproduct of the computer’s “track-then-guess” heuristic, in which it attempts to follow the player in the early and middle stages of the trial and then randomly guesses a direction to move late in play. As a result, during the early and middle phases of the trial, the puck and bar are closely aligned horizontally and expected values hover near 50%, Later, after a “point of no return” at which the computer makes its last decision, expected values are bimodal and concentrated around 0 and 1, reflecting a nearly deterministic outcome.

In fact, this trend can also be visualized in terms of the density of value as a function of time in trial (Fig 7), Against the human opponent (Fig 7A), values start out concentrated around a player’s mean win rate and evolve gradually over the course of the trial toward the 0 and 1 outcomes. By contrast, against the computer, values hold around 0.5 until abruptly diverging at the critical point. And indeed, this pattern holds in the average across all participants (Fig 7C,D). Note that here, in the case of a computer opponent defined by a simple heuristic, our model is easily able to recover strong indications of that heuristic in an unbiased way. This indicates that our approach is powerful enough to characterize a wide range of behavior. Circumstantially, it also suggests that our participants are unlikely to have relied on simple heuristics alone to constructing their strategies.

**Figure 7:**
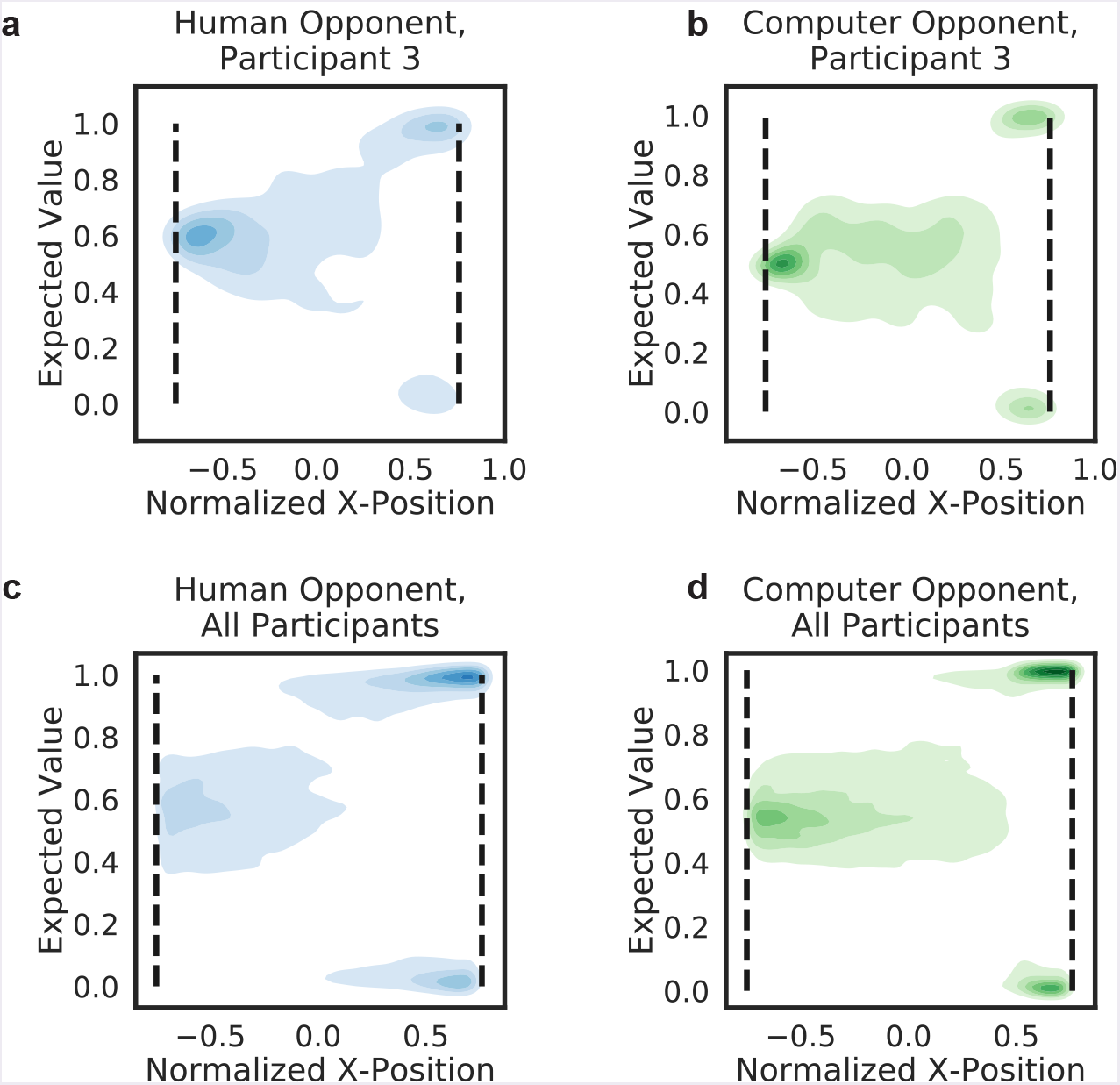
Evolution of expected value as a function of time in trial. A, B: Density of expected value as a function of time in trial for a single participant. Color indicates opponent (blue: human, green: Computer). Early in the trial, value is concentrated around the participant’s baseline win rate for each separate opponent. Over the course of the trial, values grow increasingly bimodal as the participant’s prospects for winning diverge based on game state. C, D: Average across all participants. Conventions are as in A, B. Here, the “traek-then-guess” heuristic of the computer opponent is apparent in the abrupt transition from a 50% unimodal distribution to a polarized bimodal distribution at the time of the opponent’s last move.

Finally, to investigate how well expected value predicts whether a given trial will result in a win or loss, we conducted a series of univariate logistic regressions. Given an opponent and an average expected value in the early, middle, or late periods of each trial, we attempted to predict the trial’s result, We found that regression coefficients for the human opponent condition were higher than those for the computer opponent (*t* = 4.53, *p* < 0,0001), suggesting that (unrealized) expected values better predict trial outcome in the human opponent condition. Second, we found that regression coefficients increase as the trial progresses, such that the late coefficients were significantly higher than early coefficients (*t* = 30.69, *p* < 0,0001), This matches our intuition that trial outcomes are better predicted by expected values later in the trial (see Supplement Figure 11).

## Discussion

Increasing interest in dynamic social interactions has necessitated a commensurate increase in the complexity of behavioral studies, but the methods used to analyze these new paradigms often lack the flexibility to handle the data produced. Here, we have shown that Gaussian Processes, a well-studied class of Bayesian nonparametric models, make it possible both to fit complex behavioral strategies and to forge links with the literature on reinforcement learning. As a result, our work is related to ideas in inverse reinforcement learning [44–46], which seeks to estimate, rather than learn, policies and value functions capable of generating observed behavior. Moreover, it is in keeping with a recent surge of interest in multi-agent reinforcement learning systems [47–50], though those contexts are typically cooperative rather than competitive. Finally, our problem can be viewed as a limit of the game theory context in which decisions take place simultaneously in continuous time [14,50], Our work stands to complement those results by focusing on the out-of-equilibrium dynamics that lead up to players’ final moves. This emphasis on the dynamic coupling of agents also works to bring us closer to real-world social interactions, in which decisions are based on coevolving exchanges.

There are several strengths to recommend our computational modeling framework. First, Bayesian estimation of continuous policy and value functions results in principled measures of uncertainty [28]. The resulting statistical inferences about individuals and populations are thus better indicators of model fit than point estimates obtained from maximum likelihood methods. Second, differentiability of policies and value functions allows us to derive sensitivity estimates that quantify the coupling between agents, which we have shown can characterize individual differences in play on a variety of time scales. Third, modeling the joint distribution of both players allows us to perform “counterfactual” analyses that dissociate the effects of player identity from those of game context—an intractable problem for most competing approaches [14], Finally, dissociating policy and action value functions allows us to consider observed behavior without either assuming optimality or being able to calculate what optimal behavior should be [20,51].

Importantly, our approach is not limited to a specific task. It generalizes readily to more than two agents, both cooperative and competitive contexts, and a wide variety of reward structures. All of these variants can be captured by simply enlarging the state space to accommodate the additional variables characterizing each agent. Likewise, our data need not have been sampled densely or even at regular intervals, since Gaussian Processes have proven hugely influential in fields like ecology [36] and health data [41] where sparse observations are the norm. But our method is likely to prove most valuable for examinations of decision making in natural settings like shopping, foraging, or web browsing, where the number of covariates is large and the number of events (purchases, food items, clicks) is comparatively small.

Yet our specific application does yield insights into humans’ dynamic strategic adjustments: We found that while participants exhibited a wide variety of behavioral strategies, most differences in play between human and computer opponents could be attributed to context effects, not opponent identity. That is, to first order, participants used the same approach against both opponents. Their resulting win rates depended primarily on how the opponent’s strategy interacted with their own. Nonetheless, opponent effects were present transiently at critical periods in each trial, during which the probability of a switch increased, as did the sensitivity of participants’ strategies to the opponent’s actions. In fact, comparing these opponent effects during the first and last half of each trial revealed a gradient of opponent coupling across our participant population (Fig 3D, Fig 4D).

The importance of these policy-derived metrics, particularly the sensitivities, is consistent with the findings of many groups that an ability to model the thoughts and intentions of another agent, particularly in competitive contexts, plays a central role in human social interaction [7,52–54]. For our task, in which within-trial dynamics are more variable than across-trial changes in strategy, an analysis of variance showed that most sensitivities were best characterized as instantaneous measures, but a few, including the baseline probability of switching and our sensitivity to opponent action metric, were relatively more trait-like, consistent with the idea that the underlying variability in our participant population is not in strategic heuristics but in the degree to which players’ actions are coupled to one another. This decomposition of variance for continuous, task-related predictors can be used in future studies for systematically determining whether a given covariate characterizes a trait-like or state-like process, which is particularly important when investigating individual differences in the social sciences.

Finally, we showed that an analysis of participants’ evolving prospects of winning easily distinguished between the “traek-then-guess” heuristic of the computer opponent and the more complex human opponents. Such an analysis allows us not only to assess the degree to which a given moment in the trial is critical to a player’s future prospects (the difference in action values should be large in those cases), but how successful players are in seizing these opportunities as they arise. In our case, we were unsurprised to find that moves made early on had only a modest effect on eventual wins: failure to exploit early opportunities did not necessarily ordain a loss. This is a result of the fact that action values are functions of *both* players’ strategies, and so situations arose in which the opponent’s strategy was so misguided that *any* move by the participant increased expected value.

Perhaps most important for studies of social and decision neuroscience, our models suggest a natural set of variables of interest at a hierarchy of temporal scales. While the policies and action values we derive offer instantaneous regressors at the tens of milliseconds resolution of electrophysiology, including EEG, MEG, and ECoG, these metrics can also be averaged at the trial and participant level for use with Í.\IH1 and PET, Providing computational frameworks for capturing complex temporal dynamics is crucial in learning and decision making [20,55,56], The key advantage of our approach lies in an ability to identify *both* behavioral tipping points (high sensitivity of policy) and reward tipping points (large differences in action value) and distinguish between the two. This is particularly crucial in the analysis of neural data, where one wishes to designate different types of cognitive events in addition to observational events (i.e. shifts in probability of winning without changes in action, or changes of mind) [57,58], Thus, taken together, our results and overall approach offer a new path to the use of more complex and naturalistic paradigms in the study and modeling of social interaction.

## Materials and methods

### Participants

This study was approved by the Institutional Review Board of Duke University Medical Center, Data from 82 healthy volunteers (age range: 18-48 years; 46 females; 37 males) were included in the behavioral analyses. All participants gave written informed consent to participate in this experiment and were informed that no deception would be used throughout the experiment, Two long-term participants played the role of the human opponent in the penalty shot task, but each participant played against only one human opponent. The human opponents were not members of the study team and had no stake in the outcome of the study apart from maximizing their own compensation.

Participants began the experiment with a 4 minute practice block followed by three experimental blocks, each approximately 12 minutes long. Participants played as many trials as they could within each 12 minute block, resulting in roughly 200 trials in total for each participant (approximately 100 trials per opponent condition). At the beginning of each trial, each participant was prompted to center the joystick in order for the next trial to begin, A centered fixation cross was then presented for a jittered amount of time, ranging from 1,0 to 7,5 seconds. Following the fixation cross, the identity of the opponent on the upcoming trial (either “Computer” or the name of the human opponent) was displayed in centered text for two seconds, Following the end of a trial, centered text displaying “WIN” or “LOSS” would appear on screen, indicating the previous trial’s outcome.

### Puck and Bar Dynamics

The puck was represented as a colored circle (of diameter 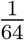 of the screen width) and started each trial at normalized coordinate position (−0.75, 0), The goal line was positioned at *x* = 0.77, The puck moved with constant horizontal velocity *υ_p_* and vertical velocity *υ_p_u_t_*, where *u_t_* ∊ [−1,1] was the vertical joystick input at time *t*. The participant controlled only the vertical velocity of the puck. The puck was constrained to remain onscreen. At each time *t*, the coordinates of the puck were updated according to:

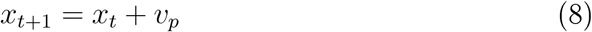

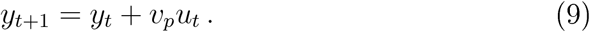

Both the human and computer opponents were identically represented on screen by a vertical bar. The bar began each trial at (0.75, 0), immediately to the left of the goal line, and could only move up or down. Unlike the puck, the opponent was able to accelerate: If the opponent maintained direction at near-maximal input (|*u*| ∊ [0.8,1]) for three consecutive time steps, the bar’s maximal velocity began to increase on the third step. That is, at each time step

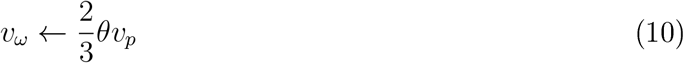

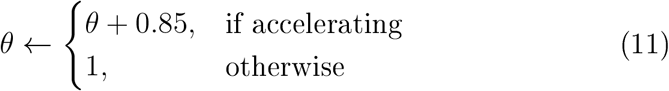

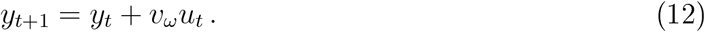

### Gaussian Process Model Fitting

Traditionally, performing full Bayesian inference in Gaussian processes has been prohibitive, with computation scaling as 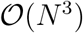, with N the number of training data points. However, recent advances in approximate inference methods based on sparse collections of *M* ≪ *N* inducing points have reduced this cost to 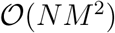, making computation feasible for large data sets [29,30,42], Here, we used GPFlow, a Gaussian process package based on the TensorFlow machine learning library, to fit separate Gaussian process classification models to data from each experimental participant [59], Models were fit using the Sparse Variational Gaussian Process algorithm coded in GPFlow, using input variables as described in the text. We used 500 inducing points and trained for 200,000 iterations using the Adam optimizer [42,59,60] for both the policy and action value models. Altering these parameters did not materially change either the fitted GPs or their sensitivities (see Supplement Figure 12,13). Model hyperparameters were learned during the training run, an empirical Bayes approach [35]. We used a train/test split of 80/20% to evaluate each model’s performance; test data were not used to select model parameters.

### Identity and Context Decomposition

Because we use a generative model to predict change points as a function of *both* game state and opponent, for any given game state, we are able to generate “counterfaetuaP predictions by providing data that were not directly observed directly in our experiment. For example, by providing the actual game state *s* for a particular moment but switching the opponent label from computer to human, we are able to predict would have happened had the participant been placed in the same game configuration against a different opponent. This allows us to assess to what degree observed differences between opponents are due to the distribution of visited game states *s* and which are due to the opponent identity ω More formally, define:

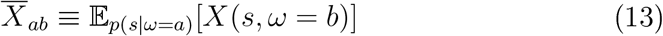

be an expectation of some random variable *x* (for instance, a probability of switching or sensitivity). Here again, *s* represents the game state and *ω* the opponent identity (0 = computer, 1 = human), but we decouple the opponent specified in the random variable from the opponent that generated the states over which we average. More concretely, *X̅*_00_ denotes the value of *X* against the computer, averaged over states actually played against the computer, while *X̅*_10_ again denotes the value of *X* against the computer, only this time averaged over states played against the *human*. In this notation, Fig 2D plots *X̅*_00_ and *X̅*_11_ with *X* = Φ^−1^(*p*), while Fig 4 shows the same two *X* equal to our opponent sensitivity metric.

What is most important, however, is that the observed contrast plotted in purple in Fig 3B-C can be decomposed as a weighted sum of the identity effect *C_identity_* and the context effect *C_identity_*, as follows:

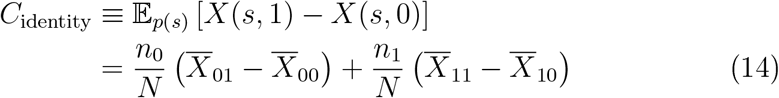

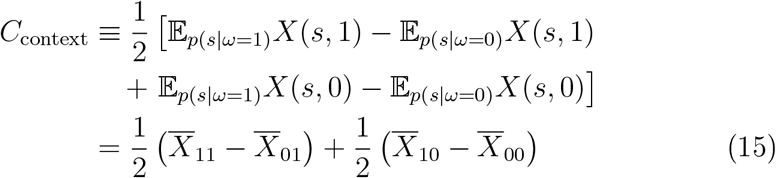

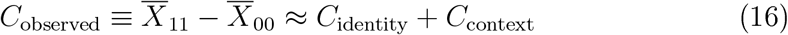

with *n*_0_ and *n_γ_* the number of trials played against the computer and human opponents, respectively, *N* = *n*_0_ + *n*_1_, and approximate equality holds in Eq 16 because *n*_0_ ≈ *n*_1_ in our data.

### Sensitivity metrics

To capture the effect of small changes of input variables on our latent Gaussian Process *f*, we a defined sensitivity for each input variable as the (squared) norm of the GP gradient along that direction:

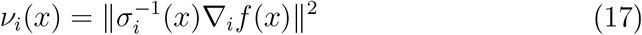

with *i* = 1 … 8 indexing each predictor variable in (*s,ω*) and *σ_i_* the local uncertainty in ∇*f*. This can be motivated by noting that since *f* is a GP, ∇*f* is as well (see Supplement Note 2), Dividing a collection of squared Gaussian variables (one per observation) by their standard deviations results in a set of *χ*^2^ variables. Viewed another way, by normalizing by the uncertainty *σ_i_*, we are downweighting highly uncertain gradients in our sensitivity measure (see Supplement Note 4).

When we consider a total sensitivity to opponent actions, we combine sensitivities to opponent action and velocity into a single metric:

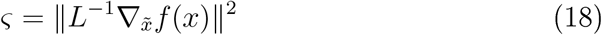

where ζ is the opponent “sensitivity” metric, *x̃* ≡ (*y_opponent_, υ_opponent_*) and *L* is the Cholesky factor of the covariance of ∇*_x̃_f* (*LL^T^* = Σ_*x*_). This is equivalent to combining the gradients for opponent position and velocity by first performing a PCA on these two coordinates and weighting each principal component equally in the calculation. As with the *v_i_* above, it can be shown that this index has a known distribution (noncentral *χ*^2^), allowing us to calculate uncertainty in the action sensitivity metric at each time point (see Supplement Note 4).

## Acknowledgments

We are thankful to the staff at Duke’s Brain Imaging Analysis Center (BIAC) for assistance with data collection as well as NIH support (S10 OD 021480) for BIAC’s super computing cluster. We are incredibly grateful to Shariq Iqbal for helpful discussions and key analysis insights. We also thank Dianna Amasino and Sam Dore for early data collection. Research reported in this publication was supported by a BD2K Career Development Award (K01-ES-025442) to JMP, an NIMH R01-108627 to SH, and a National Science Foundation Graduate Research Fellowship under NSF GRFP DGE-1644868 to KM.

## Author Contributions

W.F.B. and S.H. designed research; K.M. performed data collection; K.M. and J.P. analyzed data; and K.M., S.H., and J.P wrote the paper.

## References

[1] Von Neumann J, Morgenstern O, Theory of games and economic behavior, 2nd rev. Princeton university press; 1947.

[2] Sanfey AG, Social decision-making: insights from game theory and neuroscience. Science, 2007;318(5850):598–602.

[3] Lee D, Game theory and neural basis of social decision making. Nature neuroscience, 2008;11(4):404.

[4] Camerer CF, Behavioral game theory: Experiments in strategic interaction. Princeton University Press; 2011.

[5] Sanfey AG, Rilling JK, Aronson JA, Nystrom LE, Cohen JD, The neural basis of economic decision-making in the ultimatum game. Science, 2003;300 (5626): 1755–1758.

[6] Hampton AN, Bossaerts P, O’Doherty JP. Neural correlates of mentalizing-related computations during strategic interactions in humans. Proceedings of the National Academy of Sciences, 2008;105(18):6741–6746.

[7] Yoshida W, Dolan EJ, Friston KJ. Game theory of mind, PLoS computational biology, 2008;4(12):el000254.

[8] Yoshida W, Seymour B, Friston KJ, Dolan EJ, Neural mechanisms of belief inference during cooperative games. Journal of Neuroscience, 2010;30(32): 10744–10751.

[9] Smith JM. Price GE, The logic of animal conflict. Nature, 1973;246(5427):15.

[10] Hammerstein P, Selten E, Game theory and evolutionary biology. Handbook of game theory with economic applications, 1994;2:929–993.

[11] Platt ML, Glimcher PW, Neural correlates of decision variables in parietal cortex. Nature. 1999;400(6741):233.

[12] Eapoport A, Budescu DV, Generation of random series in two-person strictly competitive games. Journal of Experimental Psychology: General. 1992;121(3):352.

[13] Mookherjee D, Sopher B, Learning behavior in an experimental matching pennies game. Games and Economic Behavior, 1994;7(1):62–91.

[14] Delgado ME, Frank EH, Phelps EA, Perceptions of moral character modulate the neural systems of reward during the trust game. Nature neuroscience, 2005;8(11): 1611.

[15] Sally D, Conversation and cooperation in social dilemmas: A meta-analvsis of experiments from 1958 to 1992. Eationalitv and society, 1995;7(1):58–92.

[16] Zaki J, Ochsner K, The need for a cognitive neuroscience of naturalistic social cognition. Annals of the New York Academy of Sciences, 2009; 1167(1): 16–30.

[17] Iqbal S, Pearson J, A Goal-Based Movement Model for Continuous Multi-Agent Tasks. arXiv preprint arXiv: 170207319, 2017;,

[18] Kandori M, Mailath GJ, Eob E, Learning, mutation, and long run equilibria in games, Econometrica: Journal of the Econometric Society, 1993; p. 29–56.

[19] Huettel SA, Lockhead G, Psychologically rational choice: Selection between alternatives in a multiple-equilibrium game. Cognitive Systems Eesearch, 2000;1(3):143–160.

[20] Sutton ES, Barto AG, Eeinforcement learning: An introduction, vol. 1, MIT press Cambridge; 1998.

[21] Mnih V, Kavukcuoglu K, Silver D, Graves A, Antonoglou I, Wierstra D, et al. Playing atari with deep reinforcement learning, arXiv preprint arXiv:13125602, 2013;.

[22] Mnih V, Kavukcuoglu K, Silver D, Eusu AA, Veness J, Bellemare MG, et al. Human-level control through deep reinforcement learning. Nature. 2015;518(7540):529.

[23] Duryea E, Ganger M, Hu W, Exploring Deep Eeinforcement Learning with Multi Q-Learning, Intelligent Control and Automation, 2016;7(04):129.

[24] Silver D, Huang A, Maddison CJ, Guez A, Sifre L, Van Den Driessche G, et al. Mastering the game of Go with deep neural networks and tree search, nature. 2016;529(7587):484.

[25] Wang JX, Kurth-Nelson Z, Tirumala D, Soyer H, Leibo JZ, Munos E, et al. Learning to reinforcement learn, arXiv preprint arXiv:161105763, 2016XSX;.

[26] Nguyen ND, Nguyen T, Nahavandi S, System design perspective for human-level agents using deep reinforcement learning: A survey, IEEE Access. 2017;5:27091–27102.

[27] Jaderberg M, Czarnecki \Y.\I. Dunning I, Marris L, Lever G, Castaneda AG, et al. Human-level performance in first-person multiplayer games with population-based deep reinforcement learning. arXiv preprint arXiv: 180701281, 2018;.

[28] Easmussen CE, Williams CK, Gaussian process for machine learning, MIT press; 2006.

[29] Titsias M, Variational learning of inducing variables in sparse Gaussian processes. In: Artificial Intelligence and Statistics; 2009, p. 567–574.

[30] Hensman J, Fusi N, Lawrence ND. Gaussian processes for big data. arXiv preprint arXiv: 13096835. 2013;.

[31] Morya E, Eanvaud E, Pinheiro WM. Dynamics of visual feedback in a laboratory simulation of a penalty kick. Journal of Sports Sciences. 2003;21(2):87–95.

[32] Morya E, Bigatão H, Lees A, Eanvaud E. Evolving penalty kick strategies: World cup and club matches 2000-2002. Science and football V. 2005; p. 241–247.

[33] Van Der Kamp J. A field simulation study of the effectiveness of penalty kick strategies in soccer: Late alterations of kick direction increase errors and reduce accuracy. Journal of sports sciences. 2006;24(05):467–477.

[34] Dulac-Arnold G, Evans E, van Hasselt H, Sunehag P, Lillicrap T, Hunt J, et al. Deep reinforcement learning in large discrete action spaces. arXiv preprint arXiv: 151207679. 2015;.

[35] Murphy KP. Machine Learning: A Probabilistic Perspective. Adaptive Computation and Machine Learning; 2012.

[36] Gelfand AE, Diggle P, Guttorp P, Fuentes M, Handbook of spatial statistics, CEC press; 2010.

[37] Neal EM, Priors for infinite networks. In: Bayesian Learning for Neural Networks, Springer; 1996, p. 29–53.

[38] Neal EM, Bayesian learning for neural networks, vol. 118, Springer Science & Business Media; 2012.

[39] Lee J, Bahri Y, Novak E, Sehoenholz SS, Pennington J, Sohl-Diekstein J, Deep neural networks as gaussian processes, arXiv preprint arXiv:171100165, 2017;.

[40] Byron MY, Cunningham JP, Santhanam G, Evu SI, Shenov KV, Sahani M. Gaussian-process factor analysis for low-dimensional single-trial analysis of neural population activity. In: Advances in neural information processing systems; 2009. p. 1881–1888.

[41] Futoma J, Sendak M, Cameron B, Heller KA. Scalable Joint Modeling of Longitudinal and Point Process Data for Disease Trajectory Prediction and Improving Management of Chronic Kidney Disease. In: UAI; 2016.

[42] Hensman J, Matthews AG, Ghahramani Z. Scalable variational Gaussian process classification. Proceedings of Machine Learning Eesearch. 2015;38:351–360.

[43] Snoek J, Larochelle H, Adams EP, Practical bavesian optimization of machine learning algorithms. In: Advances in neural information processing systems; 2012, p, 2951–2959.

[44] Ng AY, Eussell SJ, et al. Algorithms for inverse reinforcement learning. In: Icml; 2000. p. 663–670.

[45] Abbeel P, Ng AY. Apprenticeship learning via inverse reinforcement learning. In: Proceedings of the twenty-first international conference on Machine learning. ACM; 2004. p. 1.

[46] Collette S, Pauli WM, Bossaerts P, ’Dohertv J. Neural computations underlying inverse reinforcement learning in the human brain. eLife, 2017;6. doi: 10,7554/elife.29718.

[47] Tan M, Multi-agent reinforcement learning: Independent vs, cooperative agents. In: Proceedings of the tenth international conference on machine learning; 1993, p. 330–337.

[48] Littman ML, Markov games as a framework for multi-agent reinforcement learning. In: Machine Learning Proceedings 1994, Elsevier; 1994, p. 157–163.

[49] Shoham Y, Powers R, Grenager T, Multi-agent reinforcement learning: a critical survey. Technical report, Stanford University; 2003.

[50] Braun DA, Ortega PA, Wolpert DM, Nash equilibria in multi-agent motor interactions, PLoS computational biology, 2009;5(8):el000468.

[51] Sutton ES, Me Allester DA, Singh SP, Mansour Y, Policy gradient methods for reinforcement learning with function approximation. In: Advances in neural information processing systems; 2000, p. 1057–1063.

[52] McCabe K, Houser D, Evan L, Smith V, Trouard T, A functional imaging study of cooperation in two-person reciprocal exchange. Proceedings of the National Academy of Sciences, 2001;98(20):11832–11835.

[53] Eilling JK, Sanfey AG, Aronson JA, Nystrom LE, Cohen JD, The neural correlates of theory of mind within interpersonal interactions. Neuroimage. 2004;22(4):1694–1703.

[54] Carter EM, Bowling DL, Eeeck C, Huettel SA. A distinct role of the temporal-parietal junction in predicting socially guided decisions. Science. 2012;337(6090): 109–111.

[55] Dova K. Eeinforcement learning in continuous time and space. Neural computation. 2000;12(1):219–245.

[56] Tiganj Z, Gershman SJ, Sederberg PB, Howard MW. Estimating scale-invariant future in continuous time. arXiv preprint arXiv:180206426, 2018;.

[57] Eesulaj A, Kiani E, Wolpert DM, Shadlen MN, Changes of mind in decision-making. Nature, 2009;461(7261):263.

[58] Kang YH, Petzschner FH, Wolpert DM, Shadlen MN, Piercing of consciousness as a threshold-crossing operation. Current Biology, 2017;27(15):2285–2295.

[59] Matthews AGdG, van der Wilk M, Nickson T, Fujii K, Boukouvalas A, León-Villagrá; P, et al, GPflow: A Gaussian process library using TensorFlow. Journal of Machine Learning Research, 2017;18(40): 1–6.

[60] Kingma DP, Ba J, Adam: A method for stochastic optimization, arXiv preprint arXiv: 14126980. 2014;.

